# Hydrogel mechanical properties are more important than osteoinductive growth factors for bone formation with MSC spheroids

**DOI:** 10.1101/2020.03.10.986315

**Authors:** Jacklyn Whitehead, Katherine H. Griffin, Charlotte E. Vorwald, Marissa Gionet-Gonzales, Serena E. Cinque, J. Kent Leach

**Author notes:** **Corresponding author**: J. Kent Leach, Ph.D., University of California, Davis, Department of Biomedical Engineering, Davis, CA 95616. Author contributions: JW, KHG, CEV: conception and design of experiments, data collection and assembly, data analysis and interpretation, manuscript composition MGG, SEC: conception and design of experiments, data collection and assembly, data analysis and interpretation JKL: conception and design of experiments, data analysis and interpretation, manuscript composition, administrative support.

## Abstract

Mesenchymal stromal cells (MSCs) can promote tissue repair in regenerative medicine, and their therapeutic potential is further enhanced *via* spheroid formation. We demonstrated that intraspheroidal presentation of Bone Morphogenetic Protein-2 (BMP-2) on hydroxyapatite (HA) nanoparticles resulted in more spatially uniform MSC osteodifferentiation, providing a method to internally influence spheroid phenotype. Stress relaxation of hydrogels has emerged as a potent stimulus to enhance monodispersed MSC spreading and osteogenic differentiation, but the effect of hydrogel viscoelasticity on MSC spheroids has not been reported. Herein, we describe a materials-based approach to augment the osteogenic potential of entrapped MSC spheroids by leveraging the mechanical properties of alginate hydrogels. Compared to spheroids entrapped in covalently crosslinked, elastic alginate, calcium deposition of MSC spheroids was consistently increased in ionically crosslinked, viscoelastic alginate. We observed significant increases in calcium deposition by MSC spheroids loaded with BMP-2-HA in viscoelastic gels compared to soluble BMP-2, which was higher than all elastic alginate gels. Upon implantation in critically sized calvarial bone defects, we observed enhanced bone formation in all animals treated with viscoelastic hydrogels. Increases in bone formation were evident in viscoelastic gels, regardless of the mode of presentation of BMP-2 (*i.e.*, soluble delivery or HA nanoparticles). These studies demonstrate that the dynamic mechanical properties of viscoelastic alginate are an effective strategy to enhance the therapeutic potential of MSC spheroids for bone formation and repair.

## INTRODUCTION

Mesenchymal stromal cells (MSCs) are popular in cell-based therapies because of their accessibility from several tissue compartments, trophic factor secretion, and potential to differentiate toward numerous phenotypes.[1] Among other tissues, MSCs have been used clinically to promote bone repair in the craniofacial region.[2] However, high rates of cell death upon transplantation have limited the effectiveness of such therapies [3, 4], motivating the need for alternative approaches to utilize these multipotent cells. The therapeutic efficacy of MSCs can be increased through their formation into spheroids, three-dimensional aggregates of cells.[5] Compared to monodispersed cells, spheroids exhibit improved cell survival, secrete increased concentrations of bioactive trophic factors, retain their multilineage potential, and represent a more physiologically relevant model of the native cell microenvironment.[6–8]

Osteoinductive cues such as Bone Morphogenetic Protein-2 (BMP-2) are potent stimuli to induce osteogenic differentiation of MSCs to bone-forming osteoblasts. However, soluble factors are insufficient to retain the desired cell phenotype upon the removal of these cues [5, 9, 10], as occurs upon transplantation. We demonstrated that the incorporation of hydroxyapatite (HA) nanoparticles with adsorbed BMP-2 into spheroid aggregates increased MSC osteogenic potential.[11] BMP-2-laden HA spheroids exhibited more spatially uniform expression of osteoblastic markers throughout the spheroid versus soluble BMP-2, which induced expression of these markers that was limited to the periphery of the spheroid.

The surrounding microenvironment can also influence cell phenotype, and numerous biomaterials have been studied to regulate cell function [12]. When used as a cell carrier, alginate can promote bone formation by entrapped cells due to its tunability to regulate adhesivity and substrate stiffness, along with minimally inflammatory characteristics.[13, 14] More recently, alginate was used to demonstrate the importance of viscoelasticity in a cell carrier on cell spreading and osteogenic differentiation.[15] Viscoelasticity allows entrapped cells to better engage and remodel the gel, similar to their behavior in the native extracellular matrix.[16–18] However, the role of substrate elasticity on the behavior of entrapped MSC spheroids has not been examined.

In this study, we demonstrate that the dynamic mechanical properties of viscoelastic alginate can promote the osteogenic potential of entrapped MSC spheroids. We combined an internal osteoinductive stimulus, provided by BMP-2-HA nanoparticles, with an external stimulus, alginate possessing distinct mechanical characteristics (*i.e.*, viscoelastic or elastic), to regulate the osteogenic phenotype of entrapped MSC spheroids. We then examined the capacity of MSC spheroids to promote bone formation in critical-sized calvarial defects. These data show, for the first time, that MSC spheroid behavior is dependent upon the mechanical properties of the cell carrier to guide tissue engineering strategies for intramembranous bone formation.

## MATERIALS AND METHODS

### Cell culture

Human bone marrow-derived MSCs from a 22 year old male donor (Texas A&M Institute for Regenerative Medicine, Temple, TX) were received at passage 2. We confirmed the trilineage potential of these cells as an indicator of their multipotency prior to use. MSCs were expanded in standard culture conditions (37°C, 21% O_2_, 5% CO_2_) in α-MEM (Life Technologies, Carlsbad, CA) supplemented with 10% fetal bovine serum (FBS, Atlanta Biologicals, Flowery Branch, GA) and 1% penicillin/streptomycin (P/S, Gemini Bio Products, Sacramento, CA) until use at passage 4-5.

### Spheroid formation

Spheroids were formed using a forced aggregation method.[8, 11, 19] Briefly, MSCs (4.35×10^5^ cells/mL) were pipetted into agarose molds in well plates, and the plates were centrifuged at 500x*g* for 8 min. Plates were maintained statically in standard culture conditions for 48 hr to form spheroids. Each microwell contained 15,000 MSCs.

Human recombinant BMP-2 (Medtronic, Minneapolis, MN) was adsorbed onto 0.1 mg/mL of HA nanoparticles (100 nm average diameter, Berkeley Advanced Biomaterials, Berkeley, CA) by resuspending HA in phosphate buffered saline (PBS, Thermo Fisher Scientific, Waltham, MA) as previously described.[11] In brief, BMP-2 was added to a final concentration of 200 ng/mL in low adhesion conical tubes. After 90 min, the tubes were centrifuged at 500x*g* for 8 min, and the supernatant was aspirated, leaving the pelleted HA. The pellet was washed once with PBS, the mixture centrifuged again at 500x*g* for 8 min, and PBS was aspirated. The remaining pellet was resuspended in cell culture media for subsequent incorporation into spheroids. Approximately 50 ng/mL of BMP-2 was adsorbed onto the mass of HA nanoparticles.[11]

HA was incorporated into spheroid cultures by resuspending the cell pellet of known cell number in media containing HA. The HA-cell mixture was pipetted up and down for 5 s to ensure homogeneity. The HA-cell suspension was then dispersed into the agarose molds and spheroids were formed as described above. Approximately 1 ng of BMP-2 was presented from 3 ng of HA in each spheroid.

For spheroids with soluble BMP-2 added (Sol BMP), spheroids were formed as described above except BMP-2 was not adsorbed to the HA. 50 ng/mL of BMP-2 was incorporated into the alginate hydrogel during formation.

### Alginate hydrogel formation

Arginine-Glycine-Aspartic acid (RGD)-modified alginate was prepared as described.[20–22] Briefly, G_4_RGDSP (Commonwealth Biotechnologies, Richmond, VA) was covalently coupled to UltraPure MVG (>200,000 g/mol weight average) and VLVG sodium alginate (~28,000 g/mol weight average, both from Pronova, Lysaker, Norway) using standard carbodiimide chemistry, yielding hydrogels with a bulk RGD density of 0.8 mM. The resulting RGD-alginate was sterile filtered and lyophilized for 4 days. Lyophilized alginate was reconstituted in PBS for ionically crosslinked alginate, and in 2-(N-morpholino)ethanesulfonic acid hydrate (MES) buffer for covalently crosslinked alginate, to obtain a 2.1% (w/v) solution. MSC spheroids were then entrapped in alginate and either ionically crosslinked with 50 mg/mL calcium carbonate (CaCO_3_) and 200 mg/mL of D-glucono-δ-lactone (GDL), or covalently crosslinked with 75 mM of dihydrazide (AAD)/1-hydroxybenzotriazole (HOBT) with 100 mg/mL of 1-ethyl-3- (dimethylaminopropyl) carbodiimide (EDC) (all from Sigma-Aldrich, St. Louis, MO) (**Table 1**). The reagents were mixed for 30 s using a 2-way stopcock to thoroughly blend all solutions with the cell suspension. The gel was cast between parallel glass plates with 1 mm thickness and incubated for 3 hr at 37 °C. Hydrogel disks were cut out with an 8 mm biopsy punch and placed in α-MEM with 10% FBS.

**Table 1.** List of hydrogel compositions used in these studies

| **Description** | **% Alginate** | **Alginate Weight Average** | **Resuspension Solution** | **Crosslinking Agent** | **Crosslinking Additive** |
| --- | --- | --- | --- | --- | --- |
| Ionically crosslinked<br>MVG | 2% | >75,000 g/mol | PBS | 50 mg/mL $\text{CaCO}_3$ | 200 mg/mL GDL |
| Ionically crosslinked<br>VLVG:MVG | 2% | <28,000 to >75,000 g/mol | PBS | 50 mg/mL $\text{CaCO}_3$ | 200 mg/mL GDL |
| Ionically crosslinked<br>VLVG | 2% | ~28,000 g/mol | PBS | 50 mg/mL $\text{CaCO}_3$ | 200 mg/mL GDL |
| Covalently crosslinked<br>VLVG | 2% | ~28,000 g/mol | MES buffer | 75 mM AAD/HOBT | 100 mg/mL EDC |

### Characterization of alginate hydrogel mechanical properties

The viscoelastic properties of alginate gels were measured using a Discovery HR2 Hybrid Rheometer (TA Instruments, New Castle, DE). An 8.0-mm-diameter Peltier plate geometry was used for hydrogels with a corresponding 8.0 mm diameter. An oscillatory strain sweep ranging from 0.004% to 4% strain was performed on each gel to obtain the linear viscoelastic region (LVR) before gel failure. At least 10 data points were collected for the LVR and averaged to obtain gel shear storage modulus after initial formation.[14] Stress relaxation of alginate hydrogels was measured using an Instron 3345 Compressive Testing System (Norwood, MA).[17] Hydrogels were loaded between two flat platens and compressed at 0.05 mm/sec for 18 s, then the top platen was held at a constant 10% strain for 132 s, for a total testing time of 2.5 min. Stress relaxation time was calculated by determining the time for the initial stress of the material to relax to half of that original value (τ_1/2_).[17] The stress relaxation time of alginate hydrogels was measured after initial formation and after 5 days in culture medium. Elastic moduli of alginate hydrogels gels were also tested on the Instron.[17] Gels were loaded between two flat platens and compressed at 0.05 mm/sec for 18 s. Elastic moduli were calculated from the linear portion of the slope of the stress–strain curve of hydrogels after initial formation and again after 5 days in culture medium. To determine mass loss, hydrogels were frozen overnight at −80 °C immediately after formation, lyophilized for two days, and then weighed to denote the initial dry weight. A corresponding set of hydrogels were kept in culture medium for 5 days, with the media changed every other day, then frozen at −80 °C overnight, lyophilized for two days, and weighed to denote the dry weight over time.

### Spheroid characterization

MSCs were formed into spheroids and maintained in osteogenic media consisting of α-MEM supplemented with 50 µg/mL ascorbate-2-phosphate, 10 mM β-glycerophosphate, and 100 nM dexamethasone (all from Sigma-Aldrich) for 12 days in monolayer culture. Spheroids were then formed in osteogenic media, for a total of 14 days of osteoinduction, after which media was refreshed with osteogenic media (for induction studies) or growth media (for retention studies).[11, 23] MSC spheroids were collected in passive lysis buffer (Promega, Madison, WI), and DNA content was determined using the Quant-iT PicoGreen DNA Assay Kit (Invitrogen, Carlsbad, CA). Calcium deposition was quantified using a Stanbio™ Calcium Liquid Reagent kit (Thermo Fisher). All values are presented after subtracting initial calcium levels from acellular gels and accounting for HA as a calcium source. Cell viability was assessed by a Live/Dead assay (Life Technologies).

### Evaluation of cytoskeletal structure of spheroids entrapped in alginate

At day 7 of culture, gels were fixed with 4% paraformaldehyde at 4°C overnight, washed twice with PBS, and permeabilized with 0.05% Triton-X 100 for 5 min at room temperature. Gels were stained with Alexa Fluor 488 Phalloidin solution (Thermo Fisher; 1:40 in PBS) and incubated at room temperature for 1 hr. Gels were washed once with PBS and stained with DAPI (Thermo Fisher; 1:500 in PBS) at room temperature for 10 min. Gels were washed twice with PBS and subsequently imaged using confocal microscopy (Leica TCS SP8, Wetzlar, Germany) at 40X magnification.

### Detection of YAP via Western blotting

Alginate was degraded with 5 U/mL alginate lyase (A1603; Sigma) for 45 min at 37°C. Samples were lysed and homogenized with a 30-gauge needle in RIPA buffer (Thermo Fisher Scientific), and all lysates were cleared by centrifugation. Protein concentration was evaluated with a Pierce BCA Protein Assay Kit (Thermo Fisher Scientific). HeLa cell lysate (BD Biosciences, Sparks, MD) with 1:50 dilution Primary YAP1 rabbit monoclonal antibody (mAB) (ab52771; Abcam, Cambridge, MA) and HeLa cell lysate with 1:50 Primary YAP1 (phospho S127) rabbit mAB (ab76252; Abcam) were used as positive controls for YAP1 and phosphoYAP1, respectively. HeLa cell lysate was used as a negative control. Equal amounts of protein were loaded onto a 10% Nu-PAGE Bis-Tris Gel (Invitrogen) and resolved by gel electrophoresis. Proteins were transferred using the iBlot system (Invitrogen). Membranes were blocked with blocking buffer (2.5% nonfat dry milk in TBS, Tween-20, and ultrapure H2O). Primary YAP1 rabbit monoclonal antibody (mAB) (1:5000, ab52771; Abcam), Primary YAP1 (phospho S127) rabbit mAB (1:1000, ab76252; Abcam), and Primary GAPDH rabbit mAB (1:1000, #5174; Cell Signaling Technology, Danvers, MA) were added in blocking buffer as recommended by the manufacturer. Membranes were washed with 10X TBS, Tween-20, and ultrapure H_2_O. Anti-rabbit IgG HRP-linked antibody (1:1000, #7074; Cell Signaling) was added in blocking buffer. Membranes were washed and detection was performed using the ChemiDoc MP Imaging System (BioRad, Hercules, CA).

### Assessment of implant osteogenic potential *in vivo*

Before implantation, osteogenically induced MSCs were stained with 10 µM of the near infrared (NIR) dye CellBrite NIR680 Cytoplasmic Membrane Dye (Biotium, Fremont, CA) per manufacturer’s instructions. In brief, MSCs were resuspended at 1×10^6^ in warmed culture medium containing the desired concentration of NIR dye. This cell suspension was incubated for 20 min at 37°C, protected from light. Cells were then pelleted at 500x*g* for 8 min. The supernatant was removed, and cells were washed a total of 3 times in warm medium. MSCs were aggregated into spheroids as described above. The NIR signal was detected using an IVIS Spectrum *In Vivo* Imaging System (Perkin Elmer, Waltham, MA) for fluorescence imaging (emission wavelength of 675 nm, excitation wavelength of 740 nm). Images were quantified using the IVIS’s accompanying software.

Animals were treated in accordance with all University of California, Davis animal care guidelines and National Institutes of Health (NIH) animal handling procedures. Male and female athymic rats (NIH/RNU, 10 weeks old, Taconic) were anesthetized and maintained under a 1–3% isoflurane/O_2_ mixture delivered through a nose cone. 8 mm critical-sized calvarial defects were surgically created using a trephine burr (ACE Surgical Supply, Brockton, MA). Defects were immediately filled with MSC spheroids containing BMP-2 laden HA nanoparticles entrapped in viscoelastic alginate (Viscoelastic BMP HA) or elastic alginate (Elastic BMP HA). As a control group, an equal dose of soluble BMP-2 was mixed into viscoelastic alginate gels containing HA-loaded MSC spheroids (Viscoelastic Sol BMP). The incision was closed with sutures and/or wound clips and buprenorphine (0.05 mg/kg) was administered twice per day for two days as analgesia.

### Quantification of bone formation

Two weeks post-implantation, constructs were collected, imaged for detection of NIR signal, fixed in 10% formalin for 24 h, demineralized in Calci-Clear Rapid (National Diagnostics, Atlanta, GA), processed, paraffin-embedded, and sectioned at 5 µm. Sections were stained with hematoxylin and eosin (H&E) and imaged using a Nikon Eclipse TE2000U microscope and Andor Zyla 5.5 sCMOS digital camera (Concord, MA). In order to visualize transplanted human cells undergoing osteogenic differentiation, sections underwent immunohistochemistry (IHC) using a primary antibody against DLX5 (AF6710, 1:20, R&D Systems, Minneapolis, MN), an early marker for BMP-responsive transcriptional activator needed for osteoblast differentiation.[24, 25]

At 12 weeks post-implantation, all remaining animals were euthanized, and skulls were explanted, fixed in 10% formalin for 24 h, and transferred to 70% ethanol. Constructs were imaged (70 kVp, 114 µA, 300 ms integration time, average of three images) at 15 µm resolution using a high-resolution microcomputed tomography (microCT) specimen scanner (µCT 35; Scanco Medical, Brüttisellen, Switzerland). Bone volume fraction (BVF) and bone mineral density (BMD) were determined from resulting images using the accompanying software. Explants were then demineralized in Calci-Clear Rapid (National Diagnostics), processed, paraffin-embedded, and sectioned at 5 µm thickness. Sections were stained with Masson’s Trichrome.

### Statistical analysis

Data are presented as means ± standard deviation, and all *in vitro* data represent a minimum of 3 independent experiments. Statistical analysis was performed using a one-way analysis of variance (ANOVA) with a Tukey’s multiple comparison *post hoc* test. All statistical analyses were performed using Prism 8 software (GraphPad, San Diego, CA); *p* values less than 0.05 were considered statistically significant. Significance is denoted by alphabetical letterings. Groups with significance do not share the same letters; ns denotes no significance among all groups.

## RESULTS

### Mechanical properties of alginate hydrogels are tunable

We investigated the tunability of stress relaxation in alginate hydrogels immediately after formation and over time. Though the molecular weights (MVG and VLVG) and crosslinking agents (ionic and covalent) were varied between groups, alginate hydrogels were morphologically comparable, with covalent hydrogels appearing slightly clearer while the ionic gels were more opaque due to entrapped calcium carbonate (**Fig. 1A**). All alginate hydrogels exhibited similar initial storage (**Fig. 1B**) and elastic moduli (**Fig. 1C**). The stress relaxation of the hydrogels was tunable based on molecular weight and crosslinking agent (**Fig. 1D**). As expected, covalently crosslinked alginate did not exhibit stress deformation under constant strain. We then quantified the time for the viscoelastic materials to relax to half of their initial value during a stress relaxation test (τ_1/2_) over 2.5 minutes (**Fig. 1E**). Viscoelastic VLVG alginate exhibited the fastest stress relaxation time, ~20 s. After 5 days in media, the dry mass in all alginate groups decreased slightly (**Fig. 1F**). However, all alginate hydrogels retained their initial elastic moduli (**Fig. 1G**). The stress relaxation characteristics of viscoelastic alginates were also retained, and elastic alginate did not relax (**Fig. 1H**). However, when quantified, the stress relaxation time decreased in all viscoelastic groups, except VLVG which remained similar to initial values (**Fig. 1I**).

**Figure 1.**
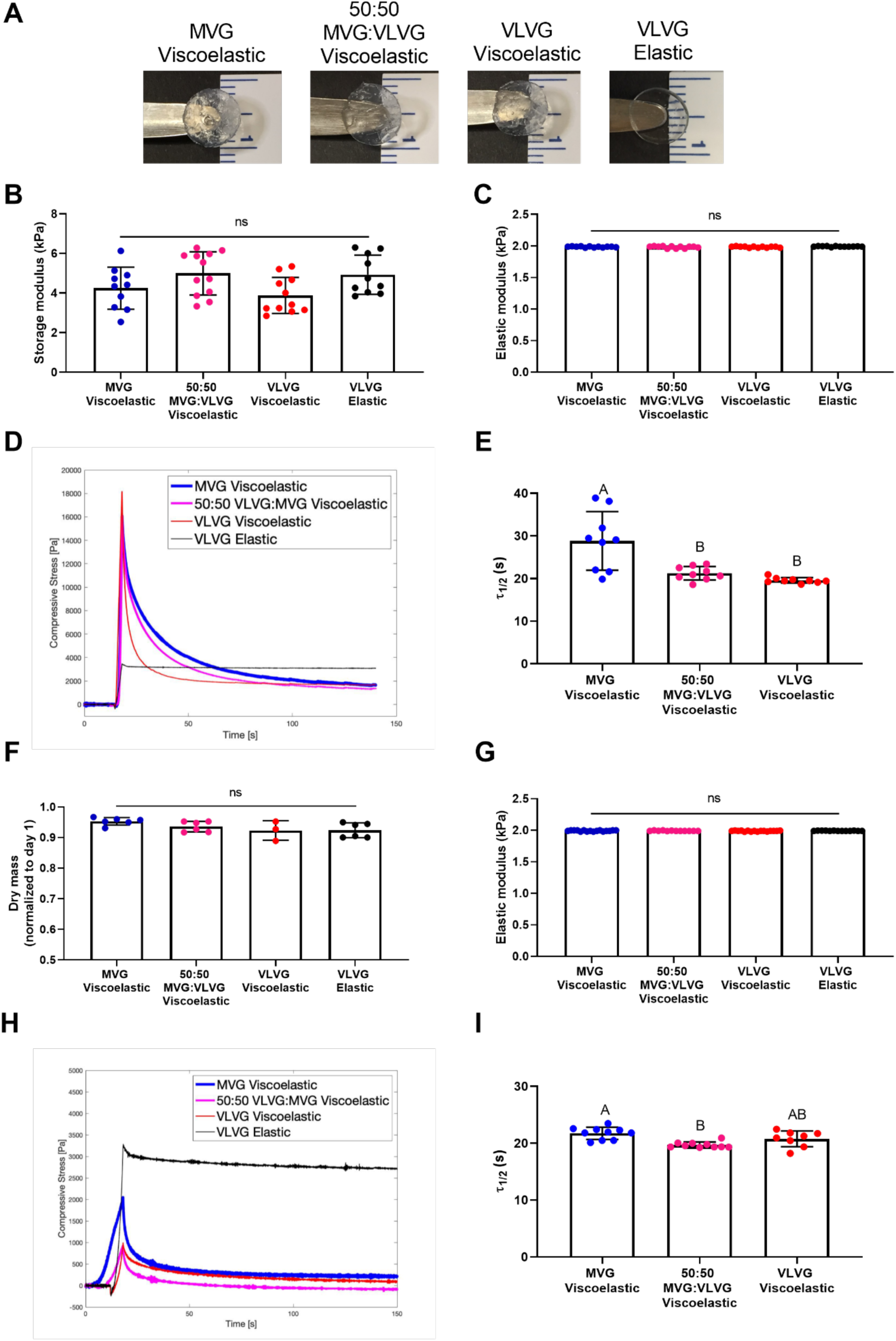
Alginate hydrogel stress relaxation is dictated by alginate molecular weight and crosslinking agent. (**A**) Representative images of alginate hydrogels immediately after formation. (**B**) Storage modulus immediately after formation as a function of molecular weight and crosslinking agent (n=10-12). (**C**) Elastic modulus as a function of molecular weight and crosslinking agent (n=12). (**D**) Representative image of the deformation of compressive stress over time as a function of crosslinking agent and alginate molecular weight. (**E**) Quantification of the stress relaxation time for alginate crosslinked with an ionic crosslinking agent as a function of molecular weight (n=9). (**F**) Quantification of dry mass of alginate hydrogels 5 days after formation normalized by the day 1 values (n=3-6). (**G**) Elastic modulus as a function of molecular weight and crosslinking agent after 5 days in cell culture medium (n=12-16). (**H**) Representative image of the deformation of compressive stress over time as a function of crosslinking agent and alginate molecular weight after 5 days in cell culture medium. (**I**) Quantification of the stress relaxation time for alginate crosslinked with an ionic crosslinking agent as a function of molecular weight after 5 days in cell culture medium (n=8-10). Significance is denoted by alphabetical letterings. Groups with significance do not share the same letters; ns denotes no significance among all groups.

### Osteogenic response of MSC spheroids is increased in viscoelastic gels

We interrogated the effect of ionically or covalently crosslinked alginate hydrogels on the osteogenic response of entrapped MSCs (**Fig. 2A**). We did not detect differences in DNA content, an indicator of cell number, when MSCs were entrapped in VLVG alginate in monodispersed or spheroidal form after 14 days (**Fig. 2B**). After 14 days in osteogenic media, MSCs in viscoelastic alginate, whether monodispersed or in spheroidal form, exhibited increased calcium deposition compared to cells in elastic gels (**Fig. 2C**).

**Figure 2.**
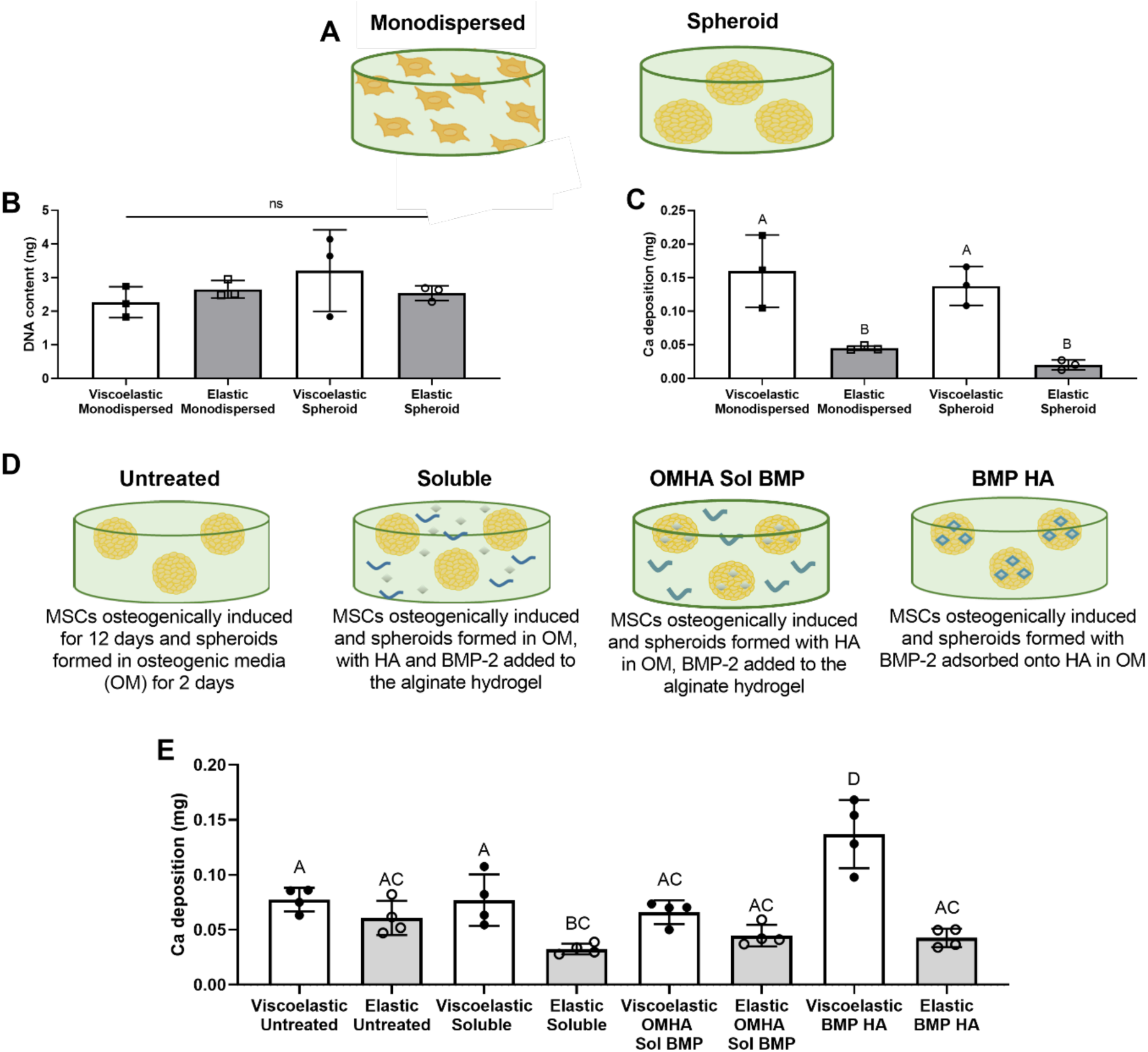
The osteogenic potential of MSC spheroids is enhanced in viscoelastic alginate. (**A**) Schematic depicting the use of MSCs in **B** and **C**. (**B**) DNA content of MSCs entrapped in viscoelastic and elastic alginate in monodispersed or spheroidal form (n=3). (**C**) Calcium deposition in gels loaded with monodispersed cells or MSC spheroids (n=3). (**D**) Schematic depicting the use of internal cues in **E**. (**E**) Calcium deposition in gels containing MSC spheroids after 14 days as a function of stimulus (n=4). Significance is denoted by alphabetical letterings. Groups with significance do not share the same letters; “ns” denotes no significance among all groups.

Having replicated the established phenomenon that monodispersed MSCs exhibit greater osteogenic potential when entrapped in viscoelastic alginate compared to elastic gels [17], the remainder of our studies used MSC spheroids to elucidate the effect of viscoelastic alginate on entrapped cellular aggregates (**Fig. 2D**). The osteogenic potential of MSC spheroids was increased when HA-loaded MSC spheroids were entrapped in viscoelastic gels (**Fig. 2E**). MSC spheroids containing BMP-2 HA nanoparticles and entrapped in viscoelastic alginate demonstrated the greatest calcium deposition compared to MSC spheroids with BMP-2 and HA in other configurations. All viscoelastic groups outperformed their elastic counterparts.

The viscoelastic characteristics of hydrogels regulate monodispersed cell spreading [15], and monodispersed cells entrapped in viscoelastic hydrogels exhibit increased nuclear localization of the transcriptional regulator yes-associated protein (YAP), a known transcriptional element mediating mechanotransduction.[26] To translate these findings to MSC spheroids, we used confocal microscopy to visualize cell spreading of MSCs on the periphery of spheroids. We also used a ROCK inhibitor, Y-27632, to restrict MSC remodeling of the viscoelastic alginate matrix (**Fig. 3A**). We observed decreased calcium deposition in viscoelastic ROCK-inhibited spheroids relative to untreated controls (*p*<0.0001) (**Fig. 3B**). Spheroids treated with soluble BMP-2 exhibited the most pronounced cell spreading in both viscoelastic and elastic hydrogels, as demonstrated by noticeably stretched actin fibers (**Fig. 3C**). When BMP-2 was delivered on HA nanoparticles to MSCs, there was little observable phalloidin staining in viscoelastic alginate. Spheroids entrapped in elastic hydrogels exhibited more pronounced phalloidin staining compared to viscoelastic hydrogels. This cell spreading was only slightly reduced upon addition of Y-27632. However, we observed substantially more YAP in spheroids entrapped in viscoelastic alginate compared to elastic alginate, regardless of the incorporation of internal stimuli (**Fig. 3D**). YAP expression was reduced with the addition of Y-27632. We detected little to no phosphorylated YAP *via* Western blot in MSC spheroids in either gel formulation (**Fig. 3E**). The presence of a strong YAP signal in MSCs entrapped in viscoelastic alginate alone is insufficient to claim osteogenic behavior. However, when combined with enhanced calcium deposition, the prominent expression of YAP that is not phosphorylated, and is thus predominantly confined to the nucleus, suggests that these hydrogels promote the osteogenic potential of MSC spheroids.

**Figure 3.**
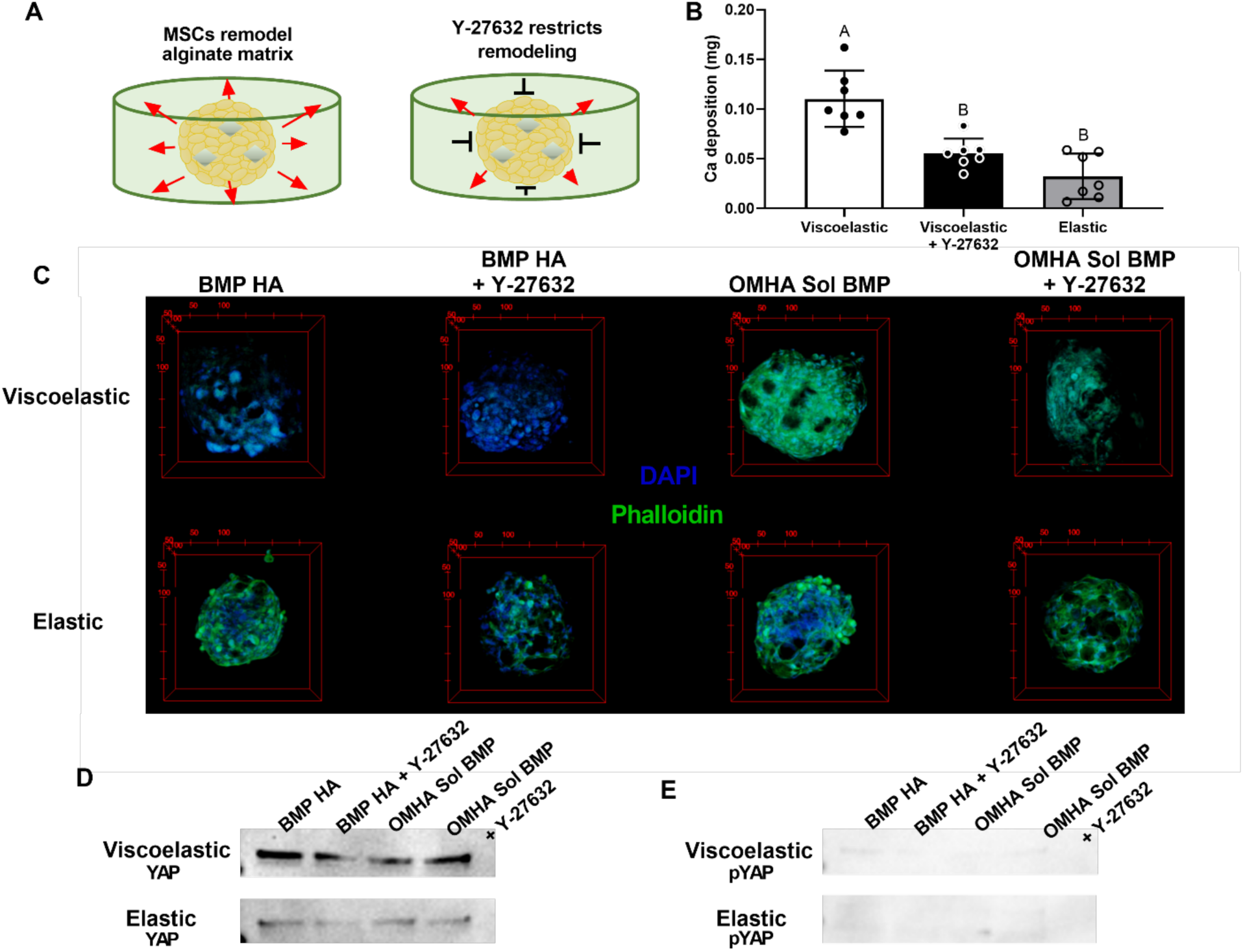
Cell spreading and YAP expression in MSC spheroids is dependent on hydrogel viscoelasticity. (**A**) Schematic representing the hypothesized influence of adding Y-27632 to alginate hydrogels. (**B**) Calcium deposition within gels containing MSC spheroids as a function of crosslinking agent and Y-27632 (n=7). Significance is denoted by alphabetical letterings. Groups with significance do not share the same letters. (**C**) Representative confocal images of the periphery of MSC spheroids with internal and external stimuli. Green is phalloidin, blue is DAPI. (**D**) Western blot for YAP as a function of internal and external stimulation of MSC spheroids. (**E**) Western blot for phosphorylated YAP as a function of internal and external stimulation of MSC spheroids.

Having established the influence of substrate mechanics and internal osteoinductive stimuli on MSC spheroid function *in vitro*, we carried forward three groups to the translational component of our study that demonstrated the greatest osteogenic potential (Viscoelastic BMP HA, Elastic BMP HA, and Viscoelastic OMHA Sol BMP). The aforementioned groups were implanted into a critical-sized defect in a rat skull, and a parallel study was performed *in vitro* to compare the translational potential of our groups. After two weeks, we examined the viability and osteogenic potential of MSC spheroids *in vitro* and *in vivo*. Spheroid viability was high in all groups (**Fig. 4A**), yet cell viability was decreased in elastic alginate hydrogels. The retained calcium deposition of both groups entrapped in viscoelastic alginate gels was significantly higher than elastic gels (**Fig. 4B**). The results from our *in vitro* characterizations were verified *in vivo* where we saw punctated DLX5 staining, an early marker for osteogenic differentiation, throughout the neotissue in all groups (**Fig. 4C**).

**Figure 4.**
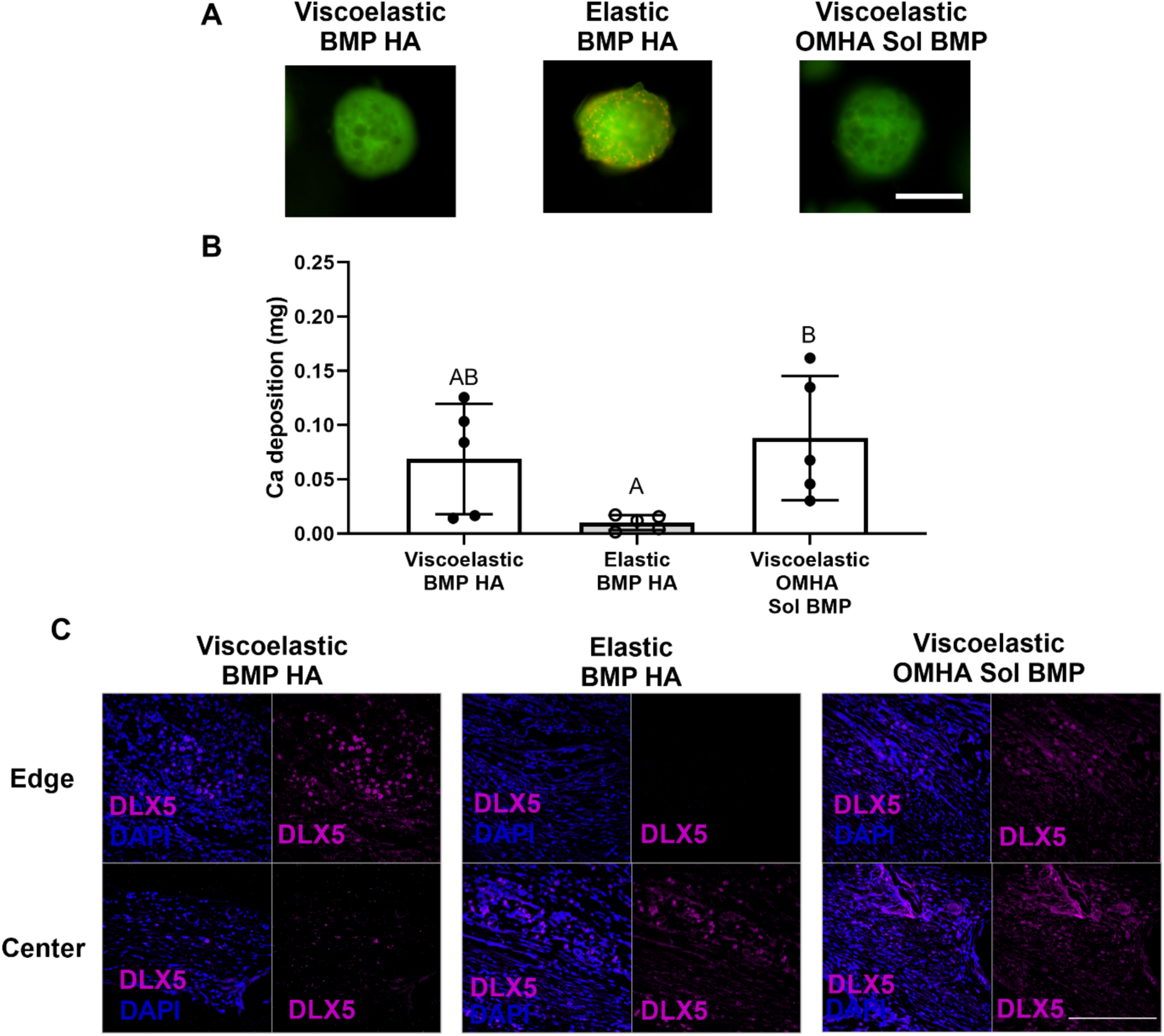
The viability and osteogenic potential of MSC spheroids is enhanced in viscoelastic alginate. (**A**) Representative images of spheroids after 14 days in culture. Live cells are green and red cells are dead. Scale bar is 500 µm. (**B**) Calcium accumulation within viscoelastic and elastic alginate hydrogels over 14 days (n=5). Significance is denoted by alphabetical letterings. Groups with significance do not share the same letters. (**C**) Representative images of DLX5 staining (magenta) counterstained with DAPI (blue) from edge or center of the calvarial defect. Scale bar is 200 µm.

We used near infrared (NIR) imaging to verify that MSC spheroids remained viable in implanted hydrogels *in vivo* two weeks after implantation. Upon explantation, MSC spheroids were visible in all calvariae, regardless of experimental group (**Fig. 5A**). The NIR signal was lower when MSCs were implanted in elastic hydrogels, yet no statistical differences were observed between groups (**Fig. 5B**). Alginate hydrogels began to dissociate after two weeks *in vivo*, as evidenced by the presence of alginate fragments in the histological sections (**Fig. 5C**).

**Figure 5.**
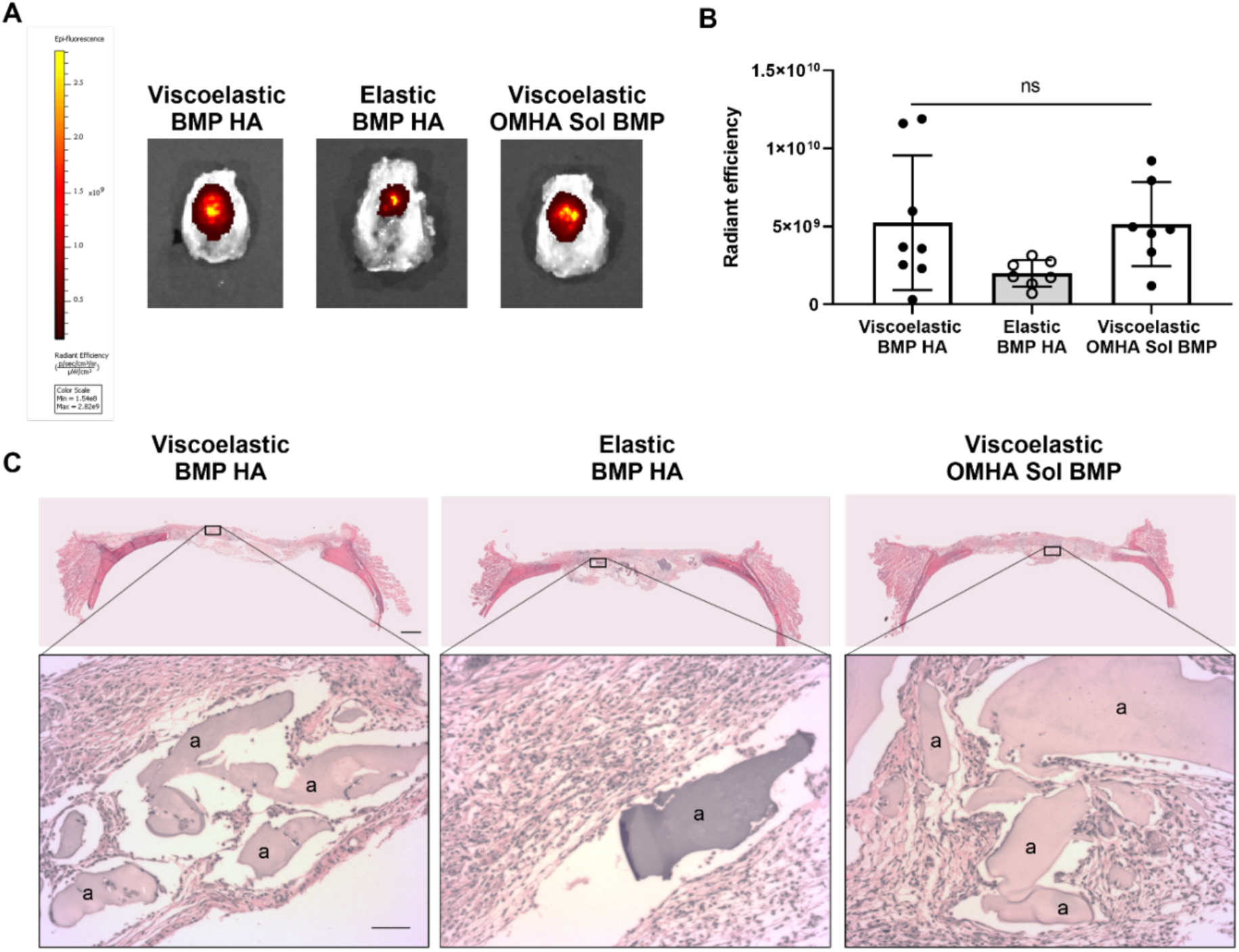
MSC spheroids persist *in vivo*. (**A**) Representative images of rat calvariae 2 weeks post-explantation exhibit NIR signal localized to the defect site. (**B**) Quantification of the NIR signal as a function of stimulus. Significance is denoted by alphabetical letterings; ns denotes no significance among all groups, n=7-8 (**C**) Representative images of H&E staining of calvarial defects from each group. Top images are large scans of an entire cross-section of the defect. Scale bar is 1 mm. The bottom insert images are 10X magnified regions of interest. The “a” denotes alginate. Bottom scale bar is 200 µm.

After 12 weeks, we observed significant differences in calcified tissue formation within treated defects using microCT (**Fig. 6A**). Defects treated with MSC spheroids entrapped in viscoelastic alginate exhibited enhanced bone volume compared to those treated with elastic alginate. However, there was no significant difference in bone volume for defects treated with MSC spheroids in viscoelastic alginate, regardless of osteoinductive stimulus (**Fig. 6B**). We observed similar trends for bone mineral density (**Fig. 6C**). Masson’s Trichrome revealed the presence of new bone formation on the edge of all defects, yet osteoid was only observed in the center of the defect with the Viscoelastic BMP HA (**Fig. 6D**).

**Figure 6.**
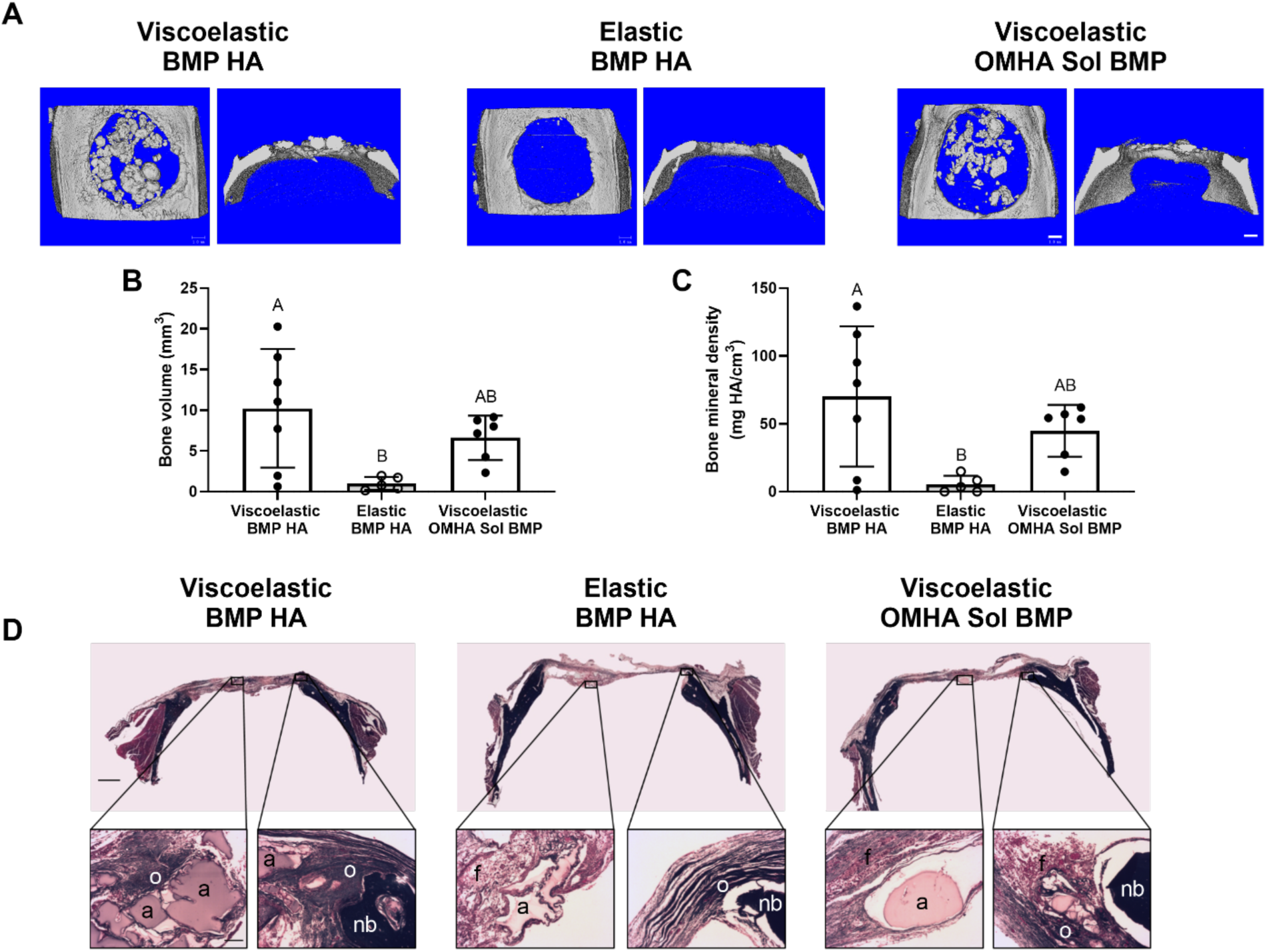
MSC spheroids promote bone tissue formation *in vivo* when entrapped in viscoelastic alginate. (**A**) Representative images of microCT scans of explanted rat calvariae 12 weeks after hydrogel implantation. Scale bars are 1 mm. (**B**) Bone volume (n=6-7). (**C**) Bone mineral density (n=6-7). Significance is denoted by alphabetical letterings. Groups with significance do not share the same letters. (**D**) Representative images of Masson’s Trichrome staining of calvarial defects from each group. Top images are large scans of an entire cross-section of the defect. Scale bar is 1 mm. The bottom insert images are 10X magnified regions of interest on either the center (left) or edge (right) of the defect. Residual alginate is denoted by “a”, fibrous tissue is denoted by “f”, native bone is denoted by “nb”, and osteoid is denoted by “o”. Bottom scale bar is 200 µm.

## DISCUSSION

Despite improved cell-cell and cell-matrix communication compared to monodispersed cells, MSC spheroids require additional instructional cues to promote their direct participation in tissue formation. In this study, we demonstrate that the dynamic mechanical properties of viscoelastic alginate can enhance *in situ* differentiation and promote the osteogenic phenotype of entrapped MSC spheroids. This work demonstrates the influence of stress relaxation properties in alginate hydrogels on the osteogenic potential of MSC spheroids. Several recent studies have reinforced the effect of substrate viscoelasticity on cell biology, namely increased cell spreading proliferation, and osteogenic differentiation.[17] However, the majority of these studies have examined the response of monodispersed cells within engineered hydrogels. Compared to monodispersed cells, we and others have demonstrated the improved therapeutic potential of MSC spheroids that possess increased survival, trophic factor secretion, and differentiation potential.[21, 27–29] In light of the promising synergy between cell spheroids and substrate viscoelasticity, this study evaluated the instructive capacity of stress relaxation on the osteogenic potential of entrapped MSC spheroids.

The stress relaxation and initial mechanical properties of alginate hydrogels are dependent upon the type and density of crosslinking agent, polymer molecular weight, and amendments to gelation protocols.[17] The interplay between cellular response and alginate mechanics has been studied under numerous conditions. For example, fibroblasts embedded in alginate gels with variable densities of polyethylene glycol (PEG), and thus variable stress relaxation characteristics, demonstrated increased cell spreading and proliferation in gels with faster stress relaxation.[30] In this study, we used an ionic crosslinking agent, CaCO_3_, to form viscoelastic alginate and a covalent crosslinking agent, AAD, to form elastic alginate. Several groups use covalent crosslinkers to produce stable and uniform gels with easily controllable mechanical properties. However, these hydrogels frequently encounter challenges during cell encapsulation that are associated with unintended and potentially adverse effects due to the crosslinking chemistry.[31, 32] Consistent with these trends, our data suggest reduced viability of MSC spheroids in the elastic hydrogel group after two weeks. These results signal the lack of bone formation observed in the same group after 12 weeks. Importantly, we observed improved cell viability of entrapped MSC spheroids compared to monodispersed cells (*data not shown*), further emphasizing the advantages of spheroids for cell-based therapies. Cell viability may be improved through the use of different covalent crosslinking agents and/or concentrations. For example, covalently crosslinked alginates formed through click chemistry of tetrazine and norbornene groups are capable of encapsulating cells without damaging them [33], while methacrylated alginates can be photocrosslinked in the presence of a photoinitiator.[34]

The influence of hydrogel stress relaxation on cell behavior has been widely reported for entrapped monodispersed cells that interact extensively with the polymer matrix. In contrast, only the cells on the periphery directly interact with the hydrogel matrix through integrin engagement, thereby motivating the need to further study the effects of dynamic mechanical properties on entrapped spheroids. MSC spheroids exhibited limited cell spreading and corresponding tension on cytoskeletal fibers when entrapped in viscoelastic alginate. Cytoskeletal tension induces translocation of YAP to the nucleus, where it activates transcriptional programs that moderate the osteogenic phenotype.[17, 26, 35] We detected increased total YAP expression, although we did not observe cell spreading of MSCs from the periphery of the spheroid in viscoelastic alginate as reported with monodispersed MSCs. Integrin engagement with the hydrogel matrix may be insufficient to stimulate cell spreading of MSCs within the spheroid, at least at the time point we investigated. Furthermore, MSCs bind to the endogenous matrix within the spheroid, as well as to one another using cadherins, both of which may limit cell spreading and migration from the aggregate. In this study, the synergistic effect of both the internal and external stimuli on MSC spheroids was sufficient to enhance YAP expression in viscoelastic alginate compared to elastic alginate. Additionally, recent studies indicate that both high cell density and three-dimensional culture decreases YAP translocation.[36, 37] Collectively, our data and the literature suggest YAP expression as an indicator of osteogenesis may be reduced compared to levels reported in conventional monolayer culture or monodispersed cells.

We previously reported that an RGD degree of substitution (DS) of 2 was sufficient to promote osteogenically induced MSC migration from spheroids.[14] However, this ligand density may have been insufficient to promote cell spreading in these studies using osteogenically induced MSCs with incorporated HA nanoparticles. Alginate hydrogels modified with higher RGD concentrations (DS=10) enhanced cell spreading of monodispersed osteogenically induced MSCs in viscoelastic alginate.[15] MSCs at later stages of osteogenic differentiation can exhibit slower migration with greater adhesivity.[38] Our previous studies demonstrate that simultaneous delivery of osteoinductive signals (BMP-2 and HA) enhanced MSC spheroid markers of osteogenic differentiation. [11] Hence, osteogenically induced MSC spheroids containing BMP-2 HA may be differentiated to a more advanced stage of osteogenesis, resulting in an inhibition of their chemotactic response to substrate-mediated cues. The role of other RGD concentrations on migration and osteogenic potential of BMP-2-HA containing MSC spheroids warrants further investigation. Additionally, the fraction of cells that could actively migrate out of the spheroid represents a small percentage of the overall cell density in the gels. Therefore, markers for the holistic behavior of the entrapped spheroids are more informative. Spheroids entrapped in viscoelastic alginate demonstrated enhanced total YAP expression and calcium deposition, both markers of the osteogenic phenotype.

This study unites several established elements in the pursuit of effective bone formation. Stress relaxation is an important stimulus to guide the proliferation and differentiation of individual or monodispersed cells, and its impact has been confirmed using numerous polymeric systems. However, compared to monodispersed cells in alginate gels, these studies are more physiologically relevant due to the increased cell-cell and cell-extracellular matrix interactions that occur within 3D cultures of cell aggregates. Studies investigating the role of these interactions in cancer progression underscore the importance of the interplay between substrate characteristics and cell behavior within dense, 3D tissue culture. For example, aggregates of mammary epithelial cells will assume malignant phenotypes when matrix stiffness is increased.[39] Our studies used human MSCs in spheroid form, which behave distinctly different than cancer cell aggregates.[40] These data establish, for the first time, the effect of stress relaxation on MSC spheroids. Consistent with findings using monodispersed MSCs, our data demonstrate that the osteogenic potential of MSC spheroids is enhanced when entrapped in viscoelastic gels. After 12 weeks *in vivo*, we observed significantly more bone tissue formation in defects treated with spheroids in viscoelastic alginate, regardless of the presence of the BMP-2. The presence of calcified material in the defect is due to an absolute regeneration of bone, evidenced by a net increase in calcified material compared to the initial mass of HA incorporated into the spheroids. Similar quantities of HA were added in both viscoelastic and elastic alginate hydrogels, yet we detected little calcified material in elastic hydrogels after 12 weeks. There was no measurable difference in calcified material between groups containing spheroids with BMP-2 HA nanoparticles or when soluble BMP-2 was added to the hydrogel. Thus, the instructive biomaterial, rather than the internal stimulus of BMP-2, advanced the osteogenic potential of entrapped MSC spheroids to a greater degree.

## CONCLUSION

These studies demonstrate that the dynamic mechanical properties of viscoelastic alginate, achieved *via* ionic crosslinking, are an effective strategy to enhance the therapeutic potential of MSC spheroids for bone formation and repair. For the first time, we demonstrated that the stress relaxation characteristics of viscoelastic alginate are retained over time and can be used for regenerative therapies using MSC spheroids. Upon entrapment in a viscoelastic hydrogel, MSC spheroids remained localized to the defect site and demonstrated heightened osteogenic potential. Thus, BMP-2 adsorbed to HA nanoparticles and incorporated into MSC spheroids, in synergy with entrapment in a viscoelastic alginate hydrogel, shows great promise for cell-based therapies to promote bone formation.

## DATA AVAILABILITY STATEMENT

The raw/processed data required to reproduce these findings will be made available on request.

## ACKNOWLEDGEMENTS

Research reported in this publication was supported by National Institute of Dental and Craniofacial Research of the National Institutes of Health under award number R01 DE025475 and R01 DE025899 to JKL. JW was supported by a National Science Foundation Graduate Research Fellowship, the Achievement Rewards for College Scientists (ARCS) Foundation fellowship, and the Schwall Dissertation Year fellowship. CEV was supported by the NHLBI Training Program in Basic and Translational Cardiovascular Science (T32 HL086350). MGG was supported by a National Science Foundation Graduate Research Fellowship. The content is solely the responsibility of the authors and does not necessarily represent the official views of the National Institutes of Health, National Science Foundation, the ARCS Foundation, or the Floyd and Mary Schwall Foundation.

## REFERENCES

[1] A.I. Caplan, D. Correa, The MSC: an injury drugstore, Cell Stem Cell 9(1) (2011) 11–15.

[2] W.L. Grayson, B.A. Bunnell, E. Martin, T. Frazier, B.P. Hung, J.M. Gimble, Stromal cells and stem cells in clinical bone regeneration, Nat Rev Endocrinol 11(3) (2015) 140–50.

[3] E. Eggenhofer, F. Luk, M.H. Dahlke, M.J. Hoogduijn, The life and fate of mesenchymal stem cells, Front Immunol 5 (2014) 148.

[4] A. Moya, J. Paquet, M. Deschepper, N. Larochette, K. Oudina, C. Denoeud, M. Bensidhoum, D. Logeart-Avramoglou, H. Petite, Human mesenchymal stem cell failure to adapt to glucose shortage and rapidly use intracellular energy reserves through glycolysis explains poor cell survival after implantation, Stem Cells 36(3) (2018) 363–376.

[5] K.C. Murphy, A.I. Hoch, J.N. Harvestine, D. Zhou, J.K. Leach, Mesenchymal stem cell spheroids retain osteogenic phenotype through α2β1 signaling, Stem Cell Transl Med 5(9) (2016) 1229–1237.

[6] Y. Xu, T.P. Shi, A.X. Xu, L. Zhang, 3D spheroid culture enhances survival and therapeutic capacities of MSCs injected into ischemic kidney, J Cell Mol Med 20(7) (2016) 1203–1213.

[7] Y. Petrenko, E. Sykova, S. Kubinova, The therapeutic potential of three-dimensional multipotent mesenchymal stromal cell spheroids, Stem Cell Res Ther 8(1) (2017) 94.

[8] C.E. Vorwald, S. Joshee, J.K. Leach, Spatial localization of endothelial cells in heterotypic spheroids influences Notch signaling, J Mol Med (Berl) (2020).

[9] A.I. Hoch, V. Mittal, D. Mitra, N. Vollmer, C.A. Zikry, J.K. Leach, Cell-secreted matrices perpetuate the bone-forming phenotype of differentiated mesenchymal stem cells, Biomaterials 74 (2016) 178–187.

[10] S. Law, S. Chaudhuri, Mesenchymal stem cell and regenerative medicine: regeneration versus immunomodulatory challenges, Am J Stem Cells 2(1) (2013) 22–38.

[11] J. Whitehead, A. Kothambawala, J.K. Leach, Morphogen delivery by osteoconductive nanoparticles instructs stromal cell spheroid phenotype, Adv Biosys (2019) 1900141.

[12] T. Gonzalez-Fernandez, P. Sikorski, J.K. Leach, Bio-instructive materials for musculoskeletal regeneration, Acta Biomater 96 (2019) 20–34.

[13] A.D. Augst, H.J. Kong, D.J. Mooney, Alginate hydrogels as biomaterials, Macromol Biosci 6(8) (2006) 623–633.

[14] S.S. Ho, A.T. Keown, B. Addison, J.K. Leach, Cell migration and bone formation from mesenchymal stem cell spheroids in alginate hydrogels are regulated by adhesive ligand density, Biomacromolecules 18(12) (2017) 4331–4340.

[15] O. Chaudhuri, L. Gu, M. Darnell, D. Klumpers, S.A. Bencherif, J.C. Weaver, N. Huebsch, D.J. Mooney, Substrate stress relaxation regulates cell spreading, Nat Commun 6 (2015) 6364.

[16] C.C. DuFort, M.J. Paszek, V.M. Weaver, Balancing forces: architectural control of mechanotransduction, Nat Rev Mol Cell Biol 12(5) (2011) 308–19.

[17] O. Chaudhuri, L. Gu, D. Klumpers, M. Darnell, S.A. Bencherif, J.C. Weaver, N. Huebsch, H.P. Lee, E. Lippens, G.N. Duda, D.J. Mooney, Hydrogels with tunable stress relaxation regulate stem cell fate and activity, Nat Mater 15(3) (2016) 326–34.

[18] X. Zhao, N. Huebsch, D.J. Mooney, Z. Suo, Stress-relaxation behavior in gels with ionic and covalent crosslinks, J Appl Phys 107(6) (2010) 63509.

[19] C.E. Vorwald, S.S. Ho, J. Whitehead, J.K. Leach, High-throughput formation of mesenchymal stem cell spheroids and entrapment in alginate hydrogels, Methods Mol Biol 1758 (2018) 139–149.

[20] J.A. Rowley, G. Madlambayan, D.J. Mooney, Alginate hydrogels as synthetic extracellular matrix materials, Biomaterials 20(1) (1999) 45–53.

[21] S.S. Ho, B.P. Hung, N. Heyrani, M.A. Lee, J.K. Leach, Hypoxic preconditioning of mesenchymal stem cells with subsequent spheroid formation accelerates repair of segmental bone defects, Stem Cells 36(9) (2018) 1393–1403.

[22] S.S. Ho, K.C. Murphy, B.Y. Binder, C.B. Vissers, J.K. Leach, Increased survival and function of mesenchymal stem cell spheroids entrapped in instructive alginate hydrogels, Stem Cells Transl Med 5(6) (2016) 773–81.

[23] A.I. Hoch, V. Mittal, D. Mitra, N. Vollmer, C.A. Zikry, J.K. Leach, Cell-secreted matrices perpetuate the bone-forming phenotype of differentiated mesenchymal stem cells, Biomaterials 74 (2016) 178–87.

[24] N. Samee, V. Geoffroy, C. Marty, C. Schiltz, M. Vieux-Rochas, G. Levi, M.C. de Vernejoul, Dlx5, a positive regulator of osteoblastogenesis, is essential for osteoblast-osteoclast coupling, Am J Pathol 173(3) (2008) 773–780.

[25] J.N. Harvestine, T. Gonzalez-Fernandez, A. Sebastian, N.R. Hum, D.C. Genetos, G.G. Loots, J.K. Leach, Osteogenic preconditioning in perfusion bioreactors improves vascularization and bone formation by human bone marrow aspirates, Science Advances 6(7) (2020) eaay2387.

[26] S. Dupont, L. Morsut, M. Aragona, E. Enzo, S. Giulitti, M. Cordenonsi, F. Zanconato, J. Le Digabel, M. Forcato, S. Bicciato, N. Elvassore, S. Piccolo, Role of YAP/TAZ in mechanotransduction, Nature 474(7350) (2011) 179–83.

[27] T.J. Bartosh, J.H. Ylostalo, A. Mohammadipoor, N. Bazhanov, K. Coble, K. Claypool, R.H. Lee, H. Choi, D.J. Prockop, Aggregation of human mesenchymal stromal cells (MSCs) into 3D spheroids enhances their antiinflammatory properties, P Natl Acad Sci USA 107(31) (2010) 13724–13729.

[28] K.C. Murphy, S.Y. Fang, J.K. Leach, Human mesenchymal stem cell spheroids in fibrin hydrogels exhibit improved cell survival and potential for bone healing, Cell Tissue Res 357(1) (2014) 91–99.

[29] P.R. Baraniak, T.C. McDevitt, Scaffold-free culture of mesenchymal stem cell spheroids in suspension preserves multilineage potential, Cell Tissue Res 347(3) (2012) 701–711.

[30] S. Nam, R. Stowers, J. Lou, Y. Xia, O. Chaudhuri, Varying PEG density to control stress relaxation in alginate-PEG hydrogels for 3D cell culture studies, Biomaterials 200 (2019) 15–24.

[31] K.Y. Lee, D.J. Mooney, Alginate: properties and biomedical applications, Prog Polym Sci 37(1) (2012) 106–126.

[32] K.A. Smeds, A. Pfister-Serres, D. Miki, K. Dastgheib, M. Inoue, D.L. Hatchell, M.W. Grinstaff, Photocrosslinkable polysaccharides for in situ hydrogel formation, J Biomed Mater Res 54(1) (2001) 115–21.

[33] R.M. Desai, S.T. Koshy, S.A. Hilderbrand, D.J. Mooney, N.S. Joshi, Versatile click alginate hydrogels crosslinked via tetrazine-norbornene chemistry, Biomaterials 50 (2015) 30–7.

[34] S.S. Ho, N.L. Vollmer, M.I. Refaat, O. Jeon, E. Alsberg, M.A. Lee, J.K. Leach, Bone morphogenetic protein-2 promotes human mesenchymal stem cell survival and resultant bone formation when entrapped in photocrosslinked alginate hydrogels, Adv Healthc Mater 5(19) (2016) 2501–2509.

[35] P.R. Nair, D. Wirtz, Enabling migration by moderation: YAP/TAZ are essential for persistent migration, J Cell Biol 218(4) (2019) 1092–1093.

[36] K. Wada, K. Itoga, T. Okano, S. Yonemura, H. Sasaki, Hippo pathway regulation by cell morphology and stress fibers, Development 138(18) (2011) 3907–14.

[37] J.Y. Lee, A.A. Dominguez, S. Nam, R.S. Stowers, L.S. Qi, O. Chaudhuri, Identification of cell context-dependent YAP-associated proteins reveals beta1 and beta4 integrin mediate YAP translocation independently of cell spreading, Sci Rep 9(1) (2019) 17188.

[38] M. Ichida, Y. Yui, K. Yoshioka, T. Tanaka, T. Wakamatsu, H. Yoshikawa, K. Itoh, Changes in cell migration of mesenchymal cells during osteogenic differentiation, FEBS Lett 585(24) (2011) 4018–24.

[39] O. Chaudhuri, S.T. Koshy, C. Branco da Cunha, J.W. Shin, C.S. Verbeke, K.H. Allison, D.J. Mooney, Extracellular matrix stiffness and composition jointly regulate the induction of malignant phenotypes in mammary epithelium, Nat Mater 13(10) (2014) 970–8.

[40] J.Y. Lee, J.K. Chang, A.A. Dominguez, H.P. Lee, S. Nam, J. Chang, S. Varma, L.S. Qi, R.B. West, O. Chaudhuri, YAP-independent mechanotransduction drives breast cancer progression, Nat Commun 10(1) (2019) 1848.

